# Faithful Supervised Dimensionality Reduction for Biomedical Data via Decision Geometry

**DOI:** 10.64898/2026.05.21.727018

**Authors:** Zexuan Wang, Zhuoping Zhou, Qipeng Zhan, Li Shen

## Abstract

Unsupervised dimensionality reduction methods aim to preserve intrinsic data geometry by maintaining local neighborhoods and approximate global relationships in low-dimensional embeddings, but they do not use label information and therefore may fail to reflect task-relevant class structure in biomedical and health applications. Supervised dimensionality reduction (SDR) incorporates labels to improve class organization, yet existing approaches often face a trade-off between discrimination and geometric faithfulness. Linear supervised methods are stable and interpretable but are limited in their ability to capture nonlinear structure, whereas many nonlinear methods impose supervision directly in the embedding space, which can over-separate classes and distort the underlying manifold. In biomedical applications, labels such as cell types in single-cell data or patient status in clinical cohorts provide meaningful biological signal, and supervised dimensionality reduction can use this information to produce more informative low-dimensional representations. Here we propose a new framework, DG-UMAP (Decision-Geometry UMAP), for faithful supervised dimensionality reduction via decision geometry. We first fit a classifier in the original feature space and use its boundary-local decision geometry to construct a low-rank metric deformation that emphasizes discriminative directions while limiting geometric distortion. Parametric UMAP is then applied to the transformed space, so supervision acts through the ambient geometry rather than by directly forcing class separation in the embedding. Across synthetic and multiple real-world biomedical datasets, our method yields embeddings with improved agreement with class structure and global organization while preserving local neighborhood quality.

## 1 Introduction

Dimensionality reduction is central to the analysis and visualization of high-dimensional data in biology, medicine and healthcare. Modern biomedical datasets—including single-cell RNA sequencing (scRNA-seq) profiles with tens of thousands of gene features, mass cytometry measurements across hundreds of protein markers, and spatial transcriptomics assays that jointly capture gene expression and tissue context, routinely involve thousands to millions of observations in spaces of very high dimension, making direct inspection intractable. By mapping observations into two or three dimensions, it can reveal structure that is otherwise difficult to inspect directly and can support interpretation, quality control and downstream modeling. Widely used unsupervised methods such as principal component analysis (PCA) [1], t-distributed stochastic neighbor embedding (t-SNE) [2], Uniform Manifold Approximation and Projection (UMAP) [3] and Potential of Heat-diffusion for Affinity-based Trajectory Embedding (PHATE) [4] emphasize different aspects of data geometry, including global variance, local neighborhood structure and progression along continuous manifolds. In many applications, however, labels or other supervision are available and carry information that is directly relevant to the scientific question. This has motivated a broad class of supervised dimensionality reduction methods, ranging from linear approaches such as linear discriminant analysis (LDA) [5] to nonlinear methods that incorporate labels through supervised graph construction [6] and metric learning approaches [3, 7].

Despite their utility, unsupervised and supervised approaches each leave an important gap. Unsupervised methods preserve intrinsic structure without using labels, and therefore may not organize the embedding according to task-relevant class relationships. As a result, classes that are separable in the original space can remain entangled in the low-dimensional representation, whereas classes that are biologically or functionally related may not be arranged in an informative way. This is particularly consequential in biomedical settings, where cell types often form hierarchical or continuous relationships—for instance, developmental trajectories in hematopoiesis or transcriptionally similar subtypes of immune cells—and where artificially collapsing or separating such structure can mislead biological interpretation. Supervised dimensionality reduction aims to address this limitation, but often does so by introducing a different problem. Linear methods are stable and interpretable, yet their global linear projections are often too restrictive for complex data. Many nonlinear supervised methods, by contrast, impose class separation directly in the embedding space. This can improve visual discrimination, but it can also exaggerate inter-class gaps, suppress within-class structure and distort the manifold geometry that unsupervised methods are designed to preserve.

To address this problem, we introduce DG-UMAP (Decision-Geometry UMAP), a supervised dimensionality reduction method that incorporates class information while preserving intrinsic data geometry. The central contribution of DG-UMAP is a different mechanism for supervision: rather than imposing class-separation objectives directly in the embedding space [6] or adding a label-related objective term [8], DG-UMAP uses decision geometry—the geometric structure induced by a classifier’s decision function in the original feature space—to reshape the ambient geometry before embedding. Specifically, DG-UMAP first fits a multiclass decision model and identifies samples near class boundaries using the margin between the top two class scores. It then extracts boundary-local score gradients, which capture directions most relevant for class separation, and aggregates them into a low-rank anisotropic deformation of the input metric. A parametric UMAP model is subsequently trained on the transformed representation using the standard attraction–repulsion objective. By introducing supervision through a decision-informed deformation of the original geometry, rather than through explicit class-forcing in the embedding, DG-UMAP aims to improve global class organization while preserving local manifold structure, avoiding artificial collapse within classes and unnecessary separation of nearby samples across different classes. These properties are especially relevant for biomedical data, where faithfully representing continuous biological variation—such as gradients of cell differentiation or disease progression—is as important as delineating discrete class boundaries.

## 2 Method

### 2.1 Problem setup

Let 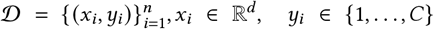,be a labeled dataset with *n* samples, ambient dimension *d*, and *c* classes. Our goal is to learn a two-dimensional embedding

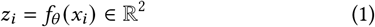

that preserves the intrinsic geometry of the data while using labels only through a controlled geometric bias, rather than through direct class-wise attraction or repulsion in the embedding. We argue that labels should not force same-class samples to collapse or different-class samples to separate, but only select a view of the data that highlights decision-relevant structure.

Linear supervised dimensionality-reduction methods impose a single global view of the data and may therefore be too restrictive when the discriminative structure is nonlinear. Nonlinear supervised methods alleviate this rigidity, but often introduce supervision by directly modifying pairwise relations. For example, supervised Isomap defines label-aware distances that contract same-class pairs and enlarge different-class pairs before computing the embedding, while supervised UMAP updates the fuzzy simplicial set using categorical label information so that neighborhood affinities more strongly reflect class membership. Such mechanisms can improve class separation, but they make labels act directly on distances or graph connectivity. By contrast, DG-UMAP uses labels only to select a decision-aware geometry, leaving the final embedding to be learned from that geometry rather than from explicit class-wise attraction or repulsion.

In reality, DG-UMAP treats supervision as a metric-selection problem for Parametric UMAP. Rather than injecting labels directly into graph affinities or the two-dimensional embedding objective, it seeks an input-space geometry that makes decision-relevant variation more visible while remaining close to the original geometry. The central object is a decision-sensitivity operator, which measures how strongly local class margins change along each direction near ambiguous samples, after balancing contributions across class pairs. DG-UMAP realizes this operator as a bounded Mahalanobis metric and then applies Parametric UMAP to the transformed data.

### 2.2 Score model and boundary measure

Let

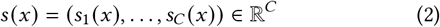

be a differentiable multiclass score model. For each sample *x*_*i*_, let

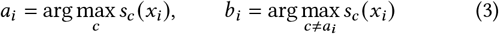

denote the highest-scoring class and its strongest competitor. We then define

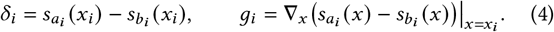

Here *δ*_*i*_ is the top-two margin gap and *g*_*i*_ is the local discriminative direction for the active class pair at *x*_*i*_ . DG-UMAP uses *δ*_*i*_ only to identify and weight ambiguous samples, and uses *g*_*i*_ later to build the decision-aware covariance.

Boundary samples are the points at which class-dependent variation is most directly expressed: they mark transitions between competing labels. By contrast, points far from such transitions mainly reflect within-class geometry and need not drive the supervised correction. We therefore localize supervision to low-margin samples, using labels to identify where the input geometry should be adjusted rather than to impose global class-wise separation. To focus on samples near class transitions, we retain the lowest *q*_*b*_ fraction of margin gaps and denote the resulting index set by ℬ. For *i* ∈ ℬ, we assign the boundary weight

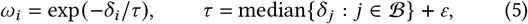

where ε *>* 0 is a small numerical constant. We then normalize these weights to obtain

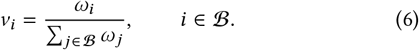

Thus *v*∈ Δ (ℬ)is a reference distribution over boundary samples that places more mass on smaller margin points. It serves as the baseline measure that is later adjusted to balance the contributions of different active class pairs.

In our experiments, we instantiate *s* as a one-vs-rest SVM, as follows.

#### Linear score model

For a one-vs-rest linear SVM,

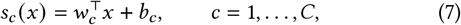

so the active-margin gradient is

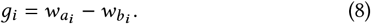

#### Exact RBF score model

For a one-vs-rest RBF SVM,

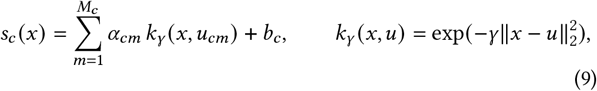

and

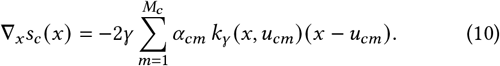

Therefore,

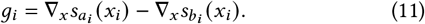

For high-dimensional inputs, we use a random Fourier feature approximation to the RBF kernel when the ambient dimension exceeds 100; see Appendix A.1.

### 2.3 Pair-balanced decision second moment

We next aggregate boundary-local directions into a positive semidefinite operator. For any sample weights *α* ∈ Δ(ℬ), define

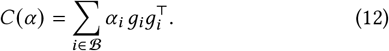

This operator averages outer products of active-margin gradients and therefore emphasizes directions along which the locally relevant class margin varies strongly near ambiguous samples.

A natural baseline is *α* = *v*. However, in multiclass data the retained set ℬ may contain many more points from some active class pairs than from others. In that case, *c* (*v*) can be dominated by pair prevalence rather than by a balanced summary of boundary structure. To reduce this effect, we separate two questions: how much total mass each active pair receives, and how that mass is distributed among the samples within that pair.

Let

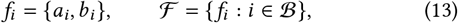

where *f*_*i*_ is the unordered active class pair at *x*_*i*_, and ℱ is the set of observed active pairs. The pair marginal induced by *v* is

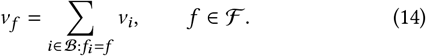

We keep the within-pair proportions from *v* and only modify the pair marginals. Concretely, for a target pair distribution *π* ∈ Δ (ℱ), we set

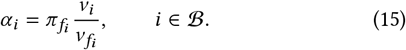

With this construction, each pair *f* receives total mass *π*_*f*_, while samples within that pair preserve the relative ordering already induced by *v*.

As a simple default, we choose *π* by interpolating between the empirical pair distribution *v*^ℱ^ = (*v*_*f*_)_*f*∈ℱ_and the uniform distribution *u* ∈ Δ(ℱ), where

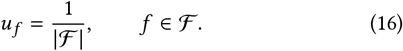

Specifically,

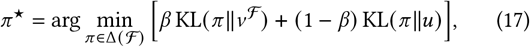

which has the closed-form solution

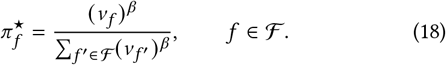

The parameter *β* ∈ [ 0, 1] controls the strength of rebalancing: *β* = 1 recovers the empirical pair frequencies, whereas *β* = 0 gives exact uniform balancing. We use 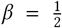 as a moderate default that reduces domination by very common pairs without forcing all pairs to contribute equally, which can over-amplify sparsely represented pairs. We do not treat 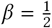 as theoretically unique; it is a practical compromise between fidelity to the empirical boundary measure and robustness to pair imbalance. The derivation is given in Appendix A.2.

Using *π*^★^, we obtain

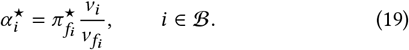

Equivalently, *α*^★^ is the information projection of *v* onto the set of sample distributions whose pair marginals equal *π*^★^:

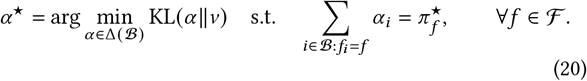

The proof is given in Appendix A.3. We then define the pair-balanced decision second moment

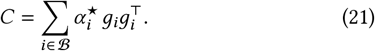

By construction, *C* is symmetric positive semidefinite. Its dominant eigenspace identifies directions along which active margins vary most strongly under the pair-balanced boundary measure, and in Appendix A.4 we show that its top-*r* eigenspace is the optimal rank-*r* subspace for capturing this weighted boundary-gradient energy.

### 2.4 A bounded low-rank decision metric

The pair-balanced decision second moment *c* identifies decision-relevant directions, but it is not yet a preprocessing metric suitable for Parametric UMAP. In particular, its scale depends on the score model and boundary weighting; it may be rank-deficient, and very small spectral components may reflect only weak evidence. To turn *C* into a practical metric, we therefore seek three additional properties: bounded distortion, full rank, and a conservative treatment of weak eigen-directions.

Let

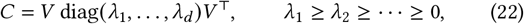

and define the relative spectral strengths

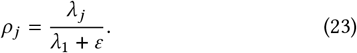

We retain

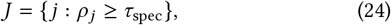

with the convention that *J* = {1} if this set is empty, and then truncate to |*J*| ≤ *r*_max_. This step keeps only directions with non-negligible evidence in *c* and prevents the metric from reacting to very small spectral components.

The framework only requires a monotone map from retained strengths *ρ*_*j*_ ∈ [0, 1] to metric gains in [1, *κ*_max_]. We use the linear map

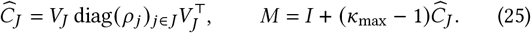

This choice preserves the ordering and relative spacing of the retained components, avoids introducing an additional shape parameter, and yields the explicit bound

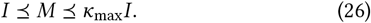

Other monotone bounded spectral maps could be used without changing the overall pipeline; we use the linear form because it is easy to interpret and keeps the construction simple.

We then transform the data by

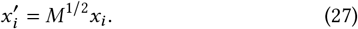

Directions orthogonal to the retained decision subspace are left unchanged, while retained directions are stretched according to their relative spectral strengths. From the bound on *M*, the preprocessing has explicit bounded distortion:

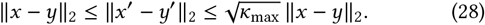

In implementation, we optionally apply a final global rescaling

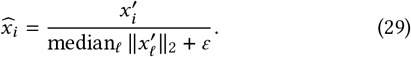

This step is only a robust numerical normalization before running Parametric UMAP. It removes an arbitrary global scale but does not change the anisotropy encoded by *M*. We use the median norm rather than the mean because it is less sensitive to outliers; any comparable global rescaling would play the same role.

#### Algorithm 1

DG-UMAP

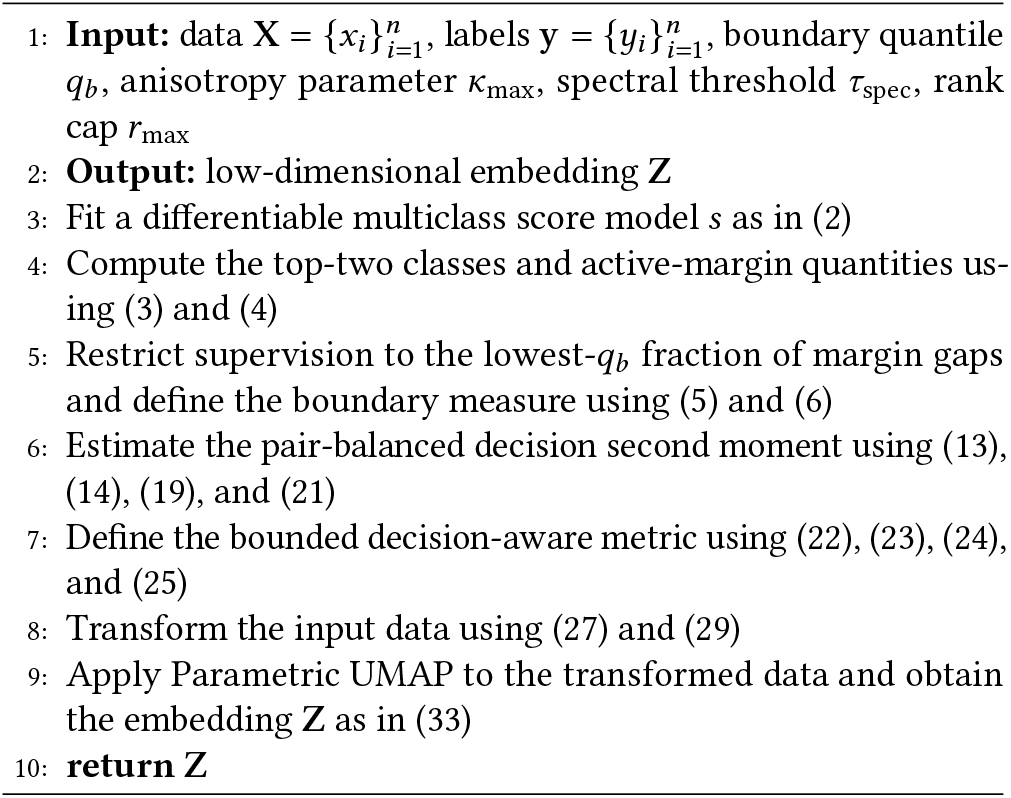

### 2.5 Parametric UMAP embedding

Given the transformed data 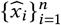, we obtain the final embedding by applying Parametric UMAP with the standard UMAP attraction– repulsion objective. UMAP first constructs a weighted fuzzy neighborhood graph

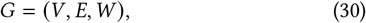

whose vertices are the samples and whose edge weights encode local neighborhood affinities under the chosen metric. It then learns a parametric encoder

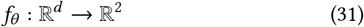

by minimizing the UMAP cross-entropy objective, so that affinities induced by distances in the two-dimensional embedding match the fuzzy neighborhood structure in the input space.

In our implementation, the encoder is a multilayer perceptron with three hidden layers of width 100 and ReLU activations, followed by a linear two-dimensional output layer, matching the architecture used in the benchmark code:

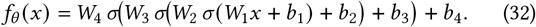

Applying this encoder to the transformed inputs yields the final two-dimensional embedding

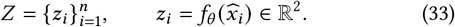

The full procedure is summarized in Algorithm 1.

## 3 Experiments

### 3.1 Experimental setup

We evaluate DG-UMAP against eleven baselines. The unsupervised baselines are UMAP [3], t-SNE [2], PHATE [4], TriMap [9], and PaCMAP [10]. The supervised baselines are S-Isomap [6], SUMAP [3], LDA [5], and NCA [7]. We also include two simple SVM-based baselines related to DG-UMAP: SVM-Score-pUMAP and SVM-WeightPCA-UMAP. SVM-Score-pUMAP applies Parametric UMAP directly to the *c*-dimensional linear SVM score vector, so supervision enters only through the classifier outputs. SVM-WeightPCA-UMAP first applies PCA to the linear SVM weight vectors, projects the input data into the resulting discriminative subspace, and then applies UMAP. Both baselines use SVM information but do not use the boundary-local, pair-balanced metric construction of DG-UMAP. For all baselines, we use either the default settings or the hyperparameter settings recommended by the original authors.

For hyperparameter settings, since DG-UMAP is a variant of the UMAP family, we used the same UMAP parameters across all experiments, with `n_neighbors=15` and `min_dist=0.05`. For t-SNE, we used `perplexity=30` and `max_iter=1000`. For PHATE, we set `k_nn=30` and `decay=40`. For PaCMAP, we used `n_neighbors=30`.

The SVM configuration follows the same settings across datasets. We trained both linear and RBF-based SVM models using `scikit-learn`. The regularization parameter was selected from

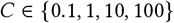

using 5-fold cross-validation. For the RBF-SVM, we used `gamma=“scale`” For the RFF-based approximation, we used `RBFSampler` with 600 random Fourier features and the same `scale` setting for the RBF bandwidth. A linear `LinearSVC` was then trained on the RFF features in a one-vs-rest manner.

DG-UMAP has three main tuned decision-geometry hyperparameters: the choice of SVM model (linear or kernel), the boundary quantile *q*_*b*_, and the maximum anisotropy *κ*_max_. Throughout our experiments, we fix the rank cap to *r*_max_ = 10 and the spectral threshold to *τ*_spec_ = 0.05.

The boundary quantile *q*_*b*_ determines which samples are used to estimate the boundary-local decision geometry. We retain the samples whose top-two margin gaps lie in the lowest *q*_*b*_ fraction. Smaller values focus the method on points very close to decision boundaries and therefore yield a more conservative correction. Larger values include a broader neighborhood around the boundaries and thus impose stronger supervision, at the risk of bringing in class-interior points. We tune *q*_*b*_ over {0.10, 0.25 } . The parameter *κ*_max_ controls the strength of the metric deformation along retained decision-relevant directions. For each retained direction *j*, the metric eigenvalue is

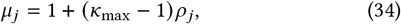

where *ρ*_*j*_ ∈ [0, 1] is the corresponding relative spectral strength. Thus *κ*_max_ sets the maximum amplification of decision-aligned directions. When *κ*_max_ is close to 1, the metric remains close to the identity, and DG-UMAP behaves similarly to standard Parametric UMAP. Larger values produce a stronger supervised deformation.

Since the corresponding axis stretch is bounded by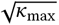, we tune *κ*_max_ over {4, 10}. The rank cap *r*_max_ limits the number of retained decision-relevant spectral directions and therefore controls the rank of the anisotropic correction. In our experiments, we fix *r*_max_ = 10, which provides a stable low-rank summary of the decision geometry without introducing an additional tuning dimension. The spectral threshold *τ*_spec_ determines which directions are retained before applying the rank cap. After computing the spectrum of the pair-balanced decision second moment, we keep only directions whose relative spectral strength exceeds *τ*_spec_. Smaller values retain more directions and yield a richer but potentially noisier metric, whereas larger values keep only the dominant directions and therefore act more aggressively as a denoising step. In our experiments, we fix *τ*_spec_ = 0.05 as a stable default.

### 3.2 Evaluation metrics

We evaluated each embedding using complementary metrics of global geometry preservation, local neighborhood fidelity, class-level organization and label recoverability. For clarity, we define the abbreviations used throughout the main text and tables as follows: global Spearman correlation (**GC**), trustworthiness (**TW**), continuity (**CT**), class energy Spearman correlation (**CEC**), and *k*-nearest-neighbor accuracy in the embedding (**kNN-Acc**) [11].

#### Global correlation (GC)

To assess preservation of large-scale structure, we computed the Spearman rank correlation between pairwise distances in the reference space and pairwise distances in the two-dimensional embedding:

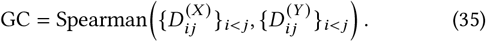

Here,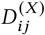 and 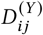 denote distances between points *i* and *j* in the reference space and embedding, respectively. For the PHATE tree, concentric cylinders and Mammoth 3D datasets, we used geodesic distances in the reference space. For MNIST and the single-cell dataset, we used Euclidean distances in the original feature space.

#### Trustworthiness (TW) and continuity (CT)

To quantify local neighborhood preservation, we used trustworthiness and continuity at neighborhood size *k*. Trustworthiness measures whether neighbors in the embedding were also neighbors in the reference space, whereas continuity measures whether neighbors in the reference space remain neighbors in the embedding. Let 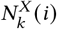 and 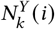 denote the *k*-nearest-neighbor sets of point *i* in the reference space and embedding, respectively. Then

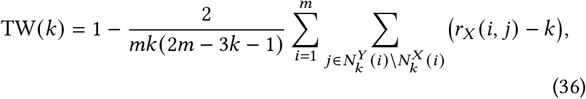

and

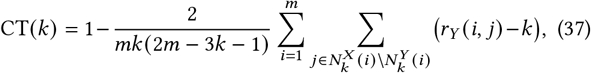

where *r*_*X*_ (*i, j*) and *r*_*Y*_ (*i, j*) are the neighbor ranks in the reference space and embedding. Higher values indicate better local fidelity. In the main text, we report TW@15 and CT@15.

#### Class energy correlation (CEC)

To evaluate preservation of class-level relationships, we compared pairwise class energy distances in the reference space and embedding. For classes *a* and *b*, the energy distance is

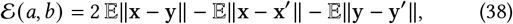

where x, x^′^ ∼*X*_*a*_ and y, y^′^∼*X*_*b*_ . We then computed the Spearman correlation between class energy distances in the reference space and embedding:

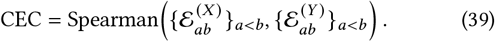

This metric reflects whether the embedding preserves the relative arrangement of class distributions.

#### k-nearest-neighbor accuracy (kNN-Acc)

As a complementary measure of label recoverability, we trained a *k*-nearest-neighbor classifier directly on the two-dimensional embedding and evaluated its accuracy on a held-out test split:

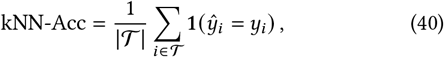

Where 𝒯 denotes the test set. This metric indicates how well class information can be recovered from the embedding, but it was interpreted together with GC, TW, CT, and CEC, since high classification accuracy alone does not imply geometric faithfulness.

We evaluated DG-UMAP on a diverse collection of synthetic and real datasets spanning image, geometric, trajectory, and single-cell settings. These datasets were selected to test whether the method can improve class organization while preserving the underlying data geometry.

#### Concentric cylinders

We generated a three-dimensional synthetic dataset consisting of three concentric cylinders that share a common central axis. Each class corresponds to one cylindrical surface with a distinct radius, while points are sampled across angular and axial coordinates and perturbed with small Gaussian noise. This dataset provides a controlled nonlinear geometry in which class identity is determined by radial structure, while continuous variation is retained along the angular and axial directions. It therefore serves as a useful benchmark for assessing whether supervised dimensionality reduction preserves both within-class geometry and global structure.

#### Mammoth 3D

We used a three-dimensional mammoth-shaped point cloud as a structured geometric benchmark [10]. The dataset contains *N* = 20, 000 points in *D* = 3 dimensions sampled from a continuous mammoth skeleton, with class labels obtained by clustering the point cloud into multiple spatial regions. Because a previous study examined preservation of inter-body-part similarity in this dataset, it provides a useful benchmark for evaluating whether a dimensionality reduction method can preserve both local and global structure.

#### High-dimensional Artificial Tree Data

We used a synthetic branching trajectory dataset generated by the PHATE framework [4], following the parameter settings of the original study. The dataset contains 3,000 points in 200 dimensions organized along a tree-structured manifold with one trunk and 10 branches of length 300, with random multiplier 2, seed 37, and Gaussian noise ( σ = 5). This dataset is useful for evaluating whether a dimensionality reduction method can reveal the global and branching structure of the data while preserving local continuity and improving branch-level organization.

#### MNIST

We used the MNIST handwritten digit dataset, which consists of grayscale images of handwritten digits represented in the original 784-dimensional pixel space [12]. MNIST is a standard benchmark for dimensionality reduction because it contains a class structure of practical interest, together with substantial within-class variability and nonlinear relationships among classes.

#### Alzheimer’s disease single-cell dataset

We evaluated DG-UMAP on a single-nucleus RNA-sequencing (snRNA-seq) dataset (ID: AD00803) obtained from the ssREAD database [13], a curated repository of AD-related single-cell transcriptomics data derived from human postmortem brain tissue. Cells are annotated with major brain cell types, including excitatory neurons, inhibitory neurons, astrocytes, oligodendrocytes, microglia, and oligodendrocyte precursor cells. Raw gene expression profiles were preprocessed and embedded into 50 principal components prior to dimensionality reduction. This dataset provides a biologically challenging setting: brain cell types in AD tissue exhibit transcriptionally continuous and partially overlapping population structure, with disease-associated transcriptional shifts further blurring boundaries between cell states. It therefore constitutes a demanding test of whether DG-UMAP can incorporate cell-type label information while preserving the continuous manifold structure characteristic of single-cell transcriptomic data.

#### Quantitative Template for the Progression of Alzheimer’s Disease Project Data

The Quantitative Template for the Progression of Alzheimer’s Disease Project Data (QT-PAD) (https://www.pi4cs.org/qt-pad-challenge) consists of heterogeneous longitudinal measurements derived from ADNI participants. As a subset of the ADNI 1/GO/2 cohort, it includes cognitive, PET, CSF, and MRI-based measures, along with covariates such as age, APOE4 status, sex, and education. QT-PAD aims to capture the ordering and timing of key biomarker and clinical changes throughout the progression to Alzheimer’s disease dementia.

#### Friedman–Nemenyi Rank Analysis Across Datasets

We assessed the consistency of method performance across datasets using a separate Friedman rank test for each evaluation metric [14]. For each method, the five repeated runs were summarized by their mean performance, and these per-dataset means were used as paired observations. Methods were ranked within each dataset, with lower ranks indicating better performance. The Friedman test evaluates whether all methods have equivalent rank distributions across datasets; a significant result indicates that at least one method differs consistently in rank. We then used the Nemenyi post-hoc test at *α* = 0.05 to compare average ranks through the critical distance. We report the average-rank profiles using radar plots and provide Friedman–Nemenyi critical-distance plots, where red points show average ranks and methods whose intervals overlap the shaded best-method region are statistically indistinguishable from the top-ranked method.

## 4 Results

Figure 1. presents qualitative embeddings produced by DG-UMAP and the benchmark methods on both synthetic and real-world datasets. Across these examples, DG-UMAP generally produces embeddings that better balance preservation of the underlying geometry with class-consistent organization. Due to space constraints, the plots for TriMap, SVM-Score-pUMAP, and SVM-WeightPCA-UMAP are included in the Appendix Figure 5. **Figure 2** summarizes the per-metric rank comparison. The left panel compares DG-UMAP with unsupervised baselines, and the right panel compares DG-UMAP with supervised baselines. These results show that DG-UMAP improves label-aware organization relative to unsupervised methods, while avoiding the stronger geometric distortion often observed in supervised baselines. The results suggest that DG-UMAP improves class-aware organization while preserving meaningful geometric structure, rather than relying on direct label forcing in the embedding space. **Figure 3** compares global geometry preservation and class-discriminative organization across datasets by plotting Global Correlation (GC) against kNN-Accuracy for each method. Across datasets, DG-UMAP lies on or near the upper-right Pareto front, indicating a favorable trade-off between geometric faithfulness and discriminative organization. **Appendix Figure 6** further reports the Friedman–Nemenyi rank analysis for the evaluated metrics. For all five metrics, the Friedman test rejects the null hypothesis of equivalent method ranks across datasets, indicating that the observed differences are consistent rather than random. In the Nemenyi plots, DG-UMAP remains within the shaded best-method comparison region for every metric, meaning that it is not significantly worse than the top-ranked method under the Nemenyi post-hoc test. DG-UMAP consistently remains among the top-performing methods across the reported metrics, indicating a stable balance between class-aware organization and geometric preservation rather than optimizing one criterion at the expense of the other.

**Figure 1:**
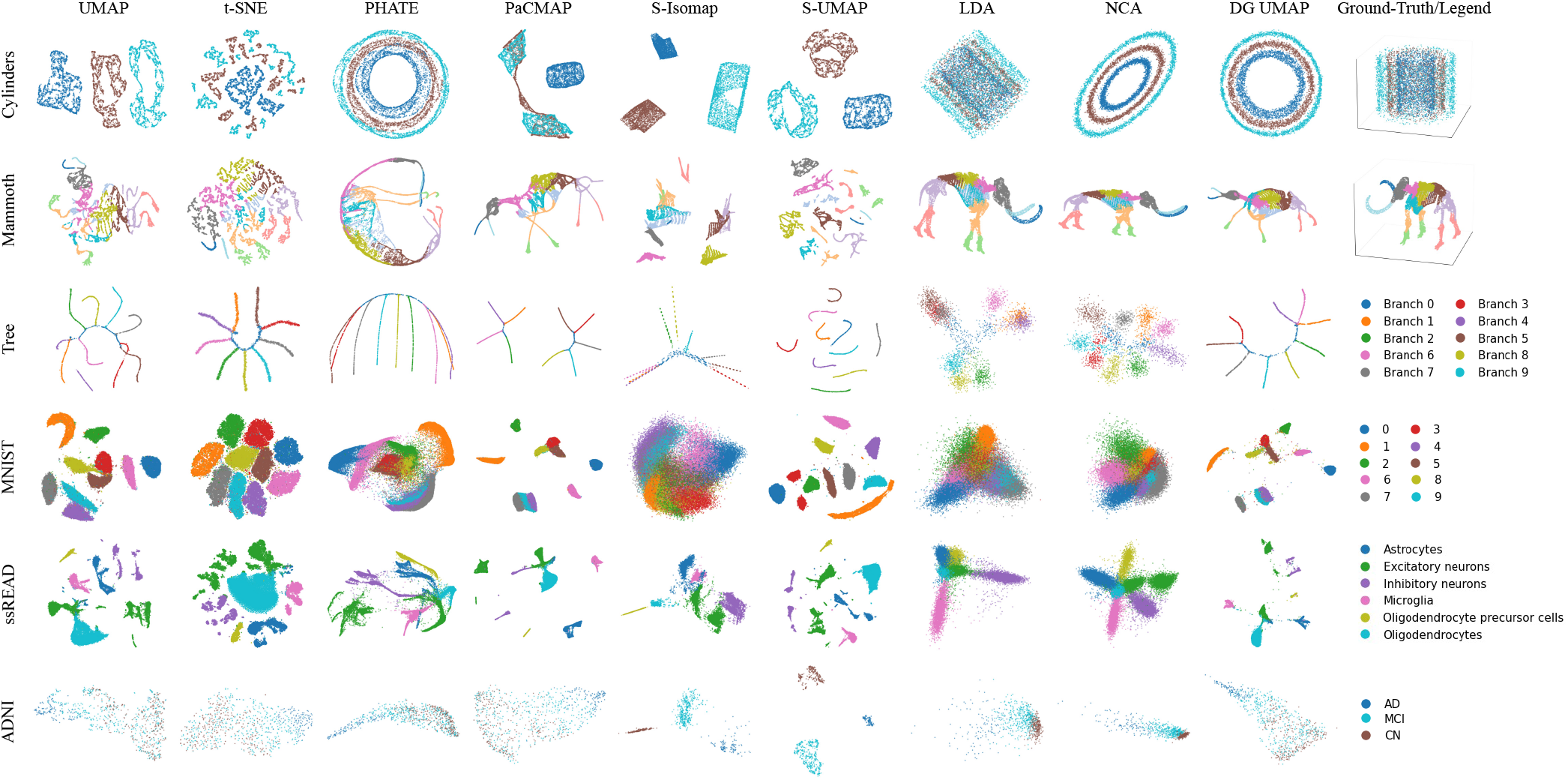
Qualitative comparison of low-dimensional embeddings produced by different dimensionality reduction methods across six datasets: `Concentric cylinders, Mammoth 3D, High-dimensional Artificial Tree Data, MNIST, ssREAD Alzheimer’s disease single-cell dataset, and ADNI QT_PAD`. Each row corresponds to one dataset, and each column shows the embedding generated by a specific method, with DG-UMAP denoting our approach. The first four methods, UMAP, t-SNE, PHATE, and PaCMAP, are unsupervised dimensionality reduction baselines and are included to show geometry-only embeddings. The next four methods, S-Isomap, S-UMAP, LDA, and NCA, are supervised dimensionality reduction baselines. The rightmost column provides the ground-truth structure or class-color legend for reference. Across datasets, DG-UMAP consistently yields embeddings that better balance preservation of intrinsic geometry and class-consistent organization than competing unsupervised and supervised baselines.

**Figure 2:**
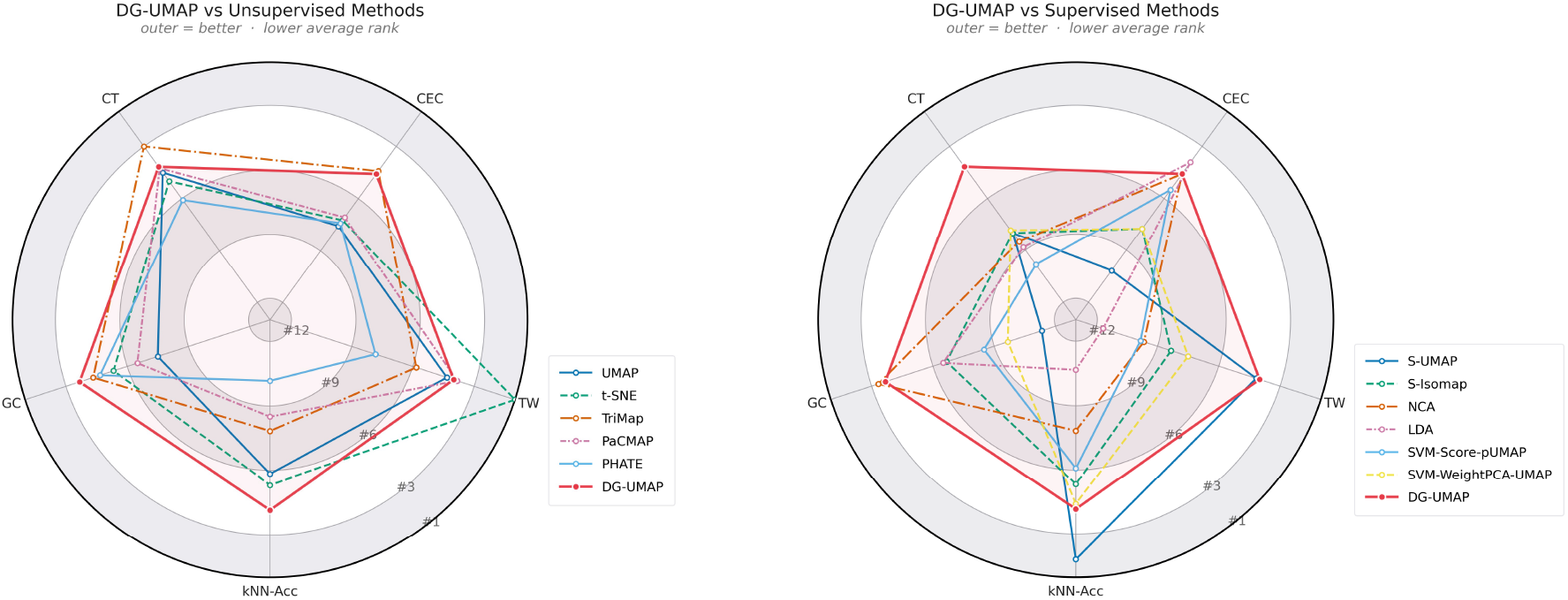
Per-metric comparison of DG-UMAP against unsupervised and supervised baselines. The left panel compares DG-UMAP with unsupervised methods, and the right panel compares DG-UMAP with supervised methods. Outer positions indicate better performance.

**Figure 3:**
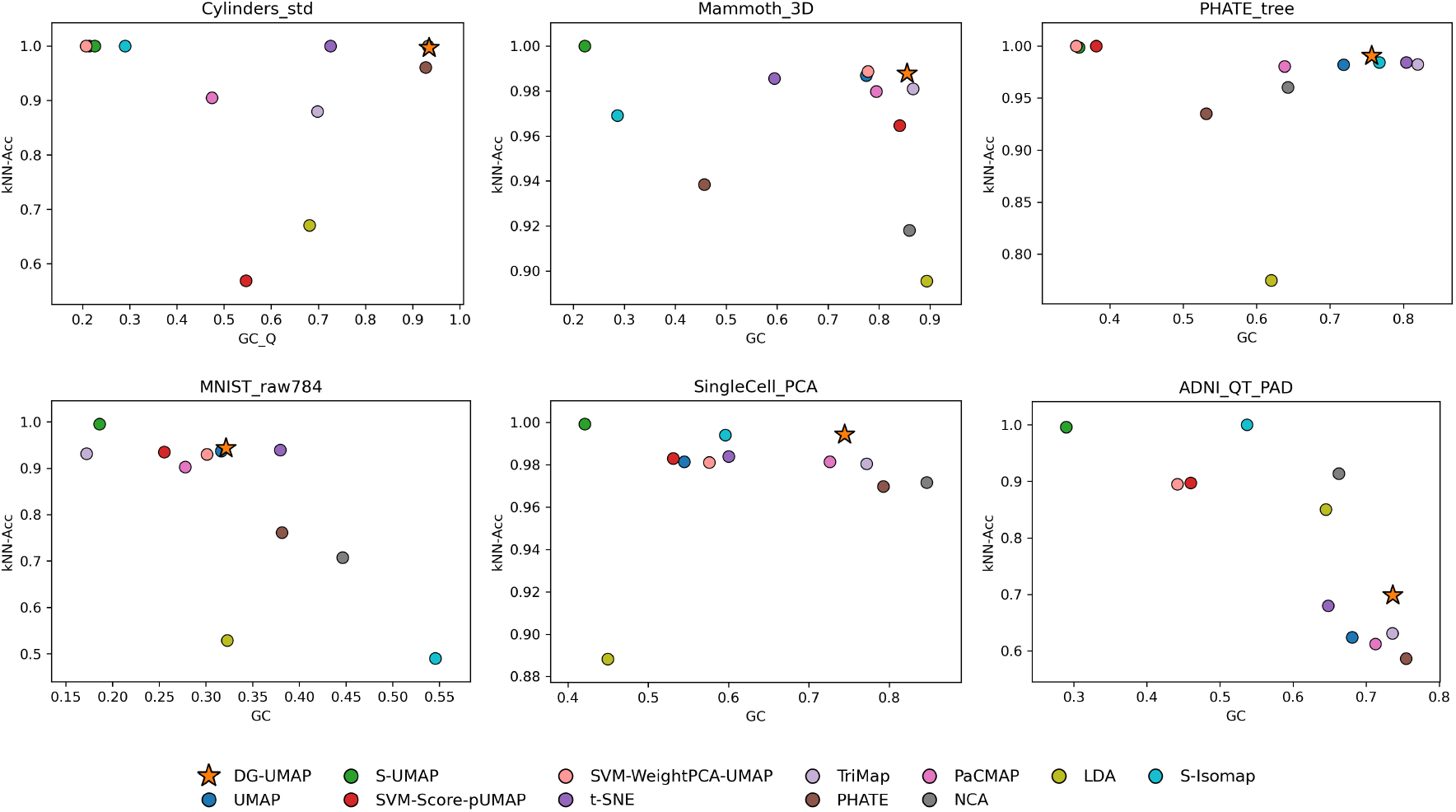
Comparison of global geometry preservation and class-discriminative organization across datasets. Each panel plots Global Correlation *Gc* against kNN-Acc for one dataset, with each point corresponding to a dimensionality reduction method and DG-UMAP highlighted by a star. Methods located toward the upper-right indicate a better trade-off between preserving global geometric structure and maintaining class-consistent neighborhood organization. Across all datasets, DG-UMAP lies on or near the Pareto front toward the upper right region, illustrating its strong overall balance between geometric fidelity and discriminative structure preservation. The remaining metrics are reported in the Appendix tables, where methods are sorted by their rank.

Appendix Tables 1–6 report the full quantitative results as mean and standard deviation over repeated runs. Appendix Table 7 reports the SVM settings and corresponding cross-validation accuracy used for each dataset.

### Concentric cylinders

On the `Cylinders` dataset, DG-UMAP gives the strongest overall result among the compared methods, as also shown by its upper-right position in Figure 3. It recovers the expected nested-ring global structure while preserving clear local organization, closely matching the ground-truth quotient view. PHATE and NCA also perform relatively well, but DG-UMAP provides the strongest overall combination of kNN-Accuracy and Global Correlation on this dataset.

By comparison, UMAP, S-UMAP, and S-Isomap preserve local neighborhoods reasonably well but distort the global concentric structure. A similar pattern is observed for the two simple DGUMAP-related baselines, SVM-Score-pUMAP and SVM-WeightPCA-UMAP. This shows that DG-UMAP better preserves both the intrinsic manifold geometry and the global class arrangement, rather than emphasizing only local separation or only coarse global shape. Methods such as t-SNE and PaCMAP fail to preserve both local neighborhood structure and global geometric arrangement. Although LDA is a supervised method, it also does not adequately recover the underlying class structure on this dataset.

### Mammoth 3D

On the `Mammoth_3D` dataset, Figure 3 shows that DG-UMAP lies on the upper-right Pareto front, indicating a strong combination of global geometry preservation and kNN-Accuracy. Consistent with this, the qualitative embedding shows that DGUMAP provides the strongest overall representation of the data manifold among the compared methods, followed by PaCMAP, TriMap, NCA, and LDA. Compared with unsupervised UMAP, which only partially recovers the global mammoth shape, DGUMAP preserves a more coherent overall structure while maintaining strong neighborhood classification performance. This suggests that the decision geometry used by DG-UMAP provides useful structural information beyond geometry alone. By contrast, the supervised baselines S-Isomap and S-UMAP tend to separate classes into disconnected groups, which weakens the underlying manifold organization.

### High-dimensional Artificial Tree Data

On the Tree dataset, Figure 3 shows that DG-UMAP lies on the Pareto front together with S-Isomap, t-SNE, and TriMap, indicating a strong combination of Global Correlation and kNN-Accuracy. However, Figure 1 shows clear differences in the resulting two-dimensional embeddings. S-Isomap produces disconnected branches, and TriMap (shown in the Appendix Figure 5) introduces intersections that do not match the true tree geometry. In contrast, DG-UMAP and t-SNE recover a reasonably coherent tree structure. PHATE also produces a relatively strong visualization, although with weaker Global Correlation. UMAP and PaCMAP introduce artificial gaps that break manifold continuity. Among the supervised baselines, LDA and NCA do not recover clear branching or finer local organization, while S-UMAP places too much emphasis on label information and distorts the relations between branches.

### MNIST

On the `MNIST` dataset, Figure 3 shows that DG-UMAP provides the second-best overall combination of Global Correlation and kNN-Accuracy. In this case, t-SNE gives the strongest overall result. DG-UMAP lies just behind t-SNE, with slightly higher kNN-Accuracy but somewhat lower Global Correlation. NCA and S-Isomap preserve Global Correlation relatively well, but this comes with a substantial drop in kNN-Accuracy. By contrast, S-UMAP places much stronger emphasis on label information, leading to very high kNN-Accuracy but very low Global Correlation.

### Alzheimer’s disease single-cell dataset

On the `Single-Cell` dataset, Figure 3 shows that DG-UMAP, NCA, and TriMap lie on the Pareto front, indicating strong overall combinations of Global Correlation and kNN-Accuracy. Among these methods, DG-UMAP provides the strongest overall embedding quality. Qualitatively, DG-UMAP produces a clearer and more coherent arrangement of the major cell populations while maintaining continuity within cell groups. By contrast, the supervised baselines S-UMAP and S-Isomap place greater emphasis on label separation, which weakens geometric faithfulness, and LDA gives the weakest overall result on this high-dimensional dataset.

### Quantitative Template for the Progression of Alzheimer’s Disease Project Data

On the `QT_PAD` dataset, Figure 3 shows that DG-UMAP lies on the Pareto front together with PHATE, NCA, and S-Isomap, indicating a strong combination of Global Correlation and kNN-Accuracy. Qualitatively, DG-UMAP preserves a clearer disease-group structure while maintaining geometric continuity. In this dataset, the expected progression is `CN→ MCI→ AD`, and most methods roughly follow this trend. However, S-UMAP tends to overemphasize label information at the expense of geometry. This is also reflected by its negative CEC value, which indicates that the overall class structure is not preserved well. Overall, these results suggest that DG-UMAP more effectively incorporates decision-guided class information while preserving the intrinsic geometry of this clinical dataset.

### Hyperparameter sensitivity

We further examined the effect of DG-UMAP hyperparameters on the `Mammoth_3D` dataset. Figure 4 shows the results in the GC–kNN-Acc plane. The upper-right cluster is produced by the linear SVM variant under different combinations of the boundary quantile *q*_*b*_ and the maximum anisotropy *κ*_max_, whereas the lower cluster is produced by the kernel SVM variant under the tested settings. For both variants, the tested configurations form relatively compact groups, indicating that the results do not vary substantially across the explored hyperparameter range.

**Figure 4:**
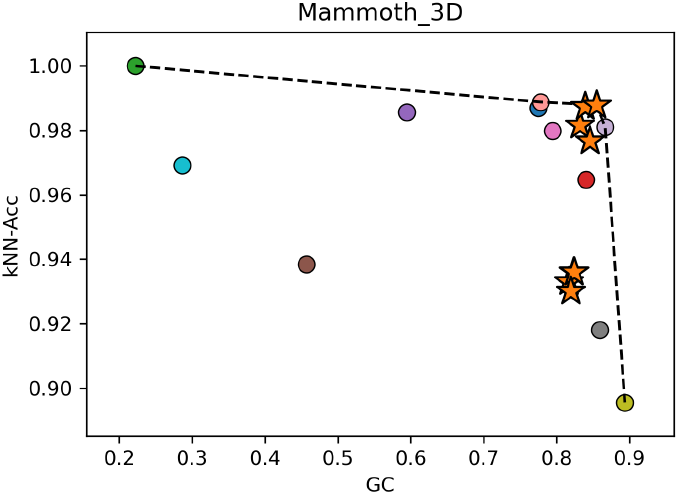
Hyperparameter sensitivity of DG-UMAP on the `Mammoth_3D` dataset, shown in the GC–kNN-Acc plane. Each star corresponds to one DG-UMAP configuration. Stars in the upper-right region are obtained with the linear SVM variant, while the lower cluster of stars corresponds to the kernel SVM variant. The dashed curve indicates the Pareto frontier, highlighting configurations that are not dominated in terms of simultaneously improving global class organization (GC) and neighborhood classification accuracy (kNN-Acc).

**Figure 5:**
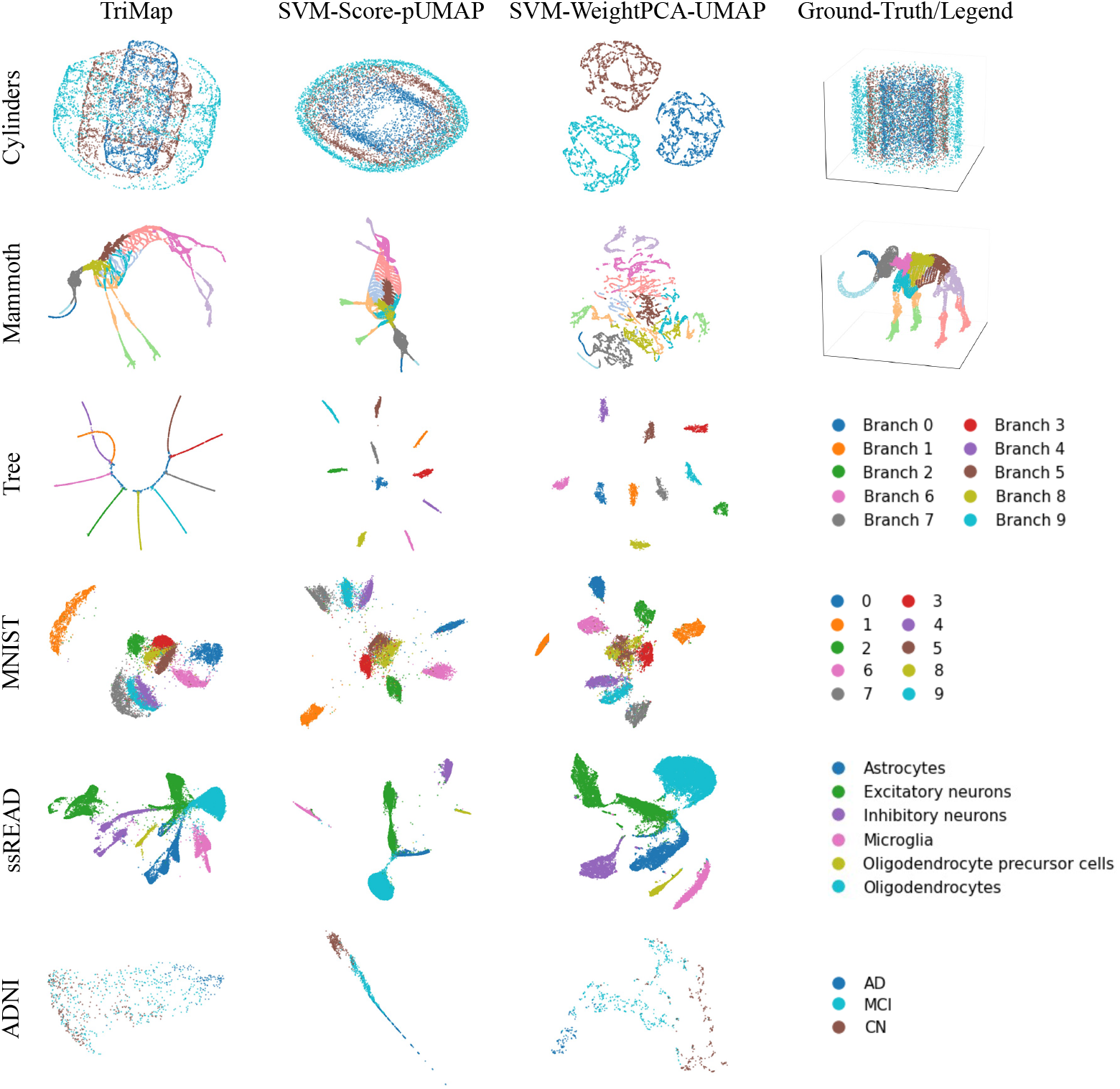
Qualitative comparison of the low-dimensional embeddings produced by TriMap, SVM-Score-pUMAP, and SVM-WeightPCA-UMAP.

**Figure 6:**
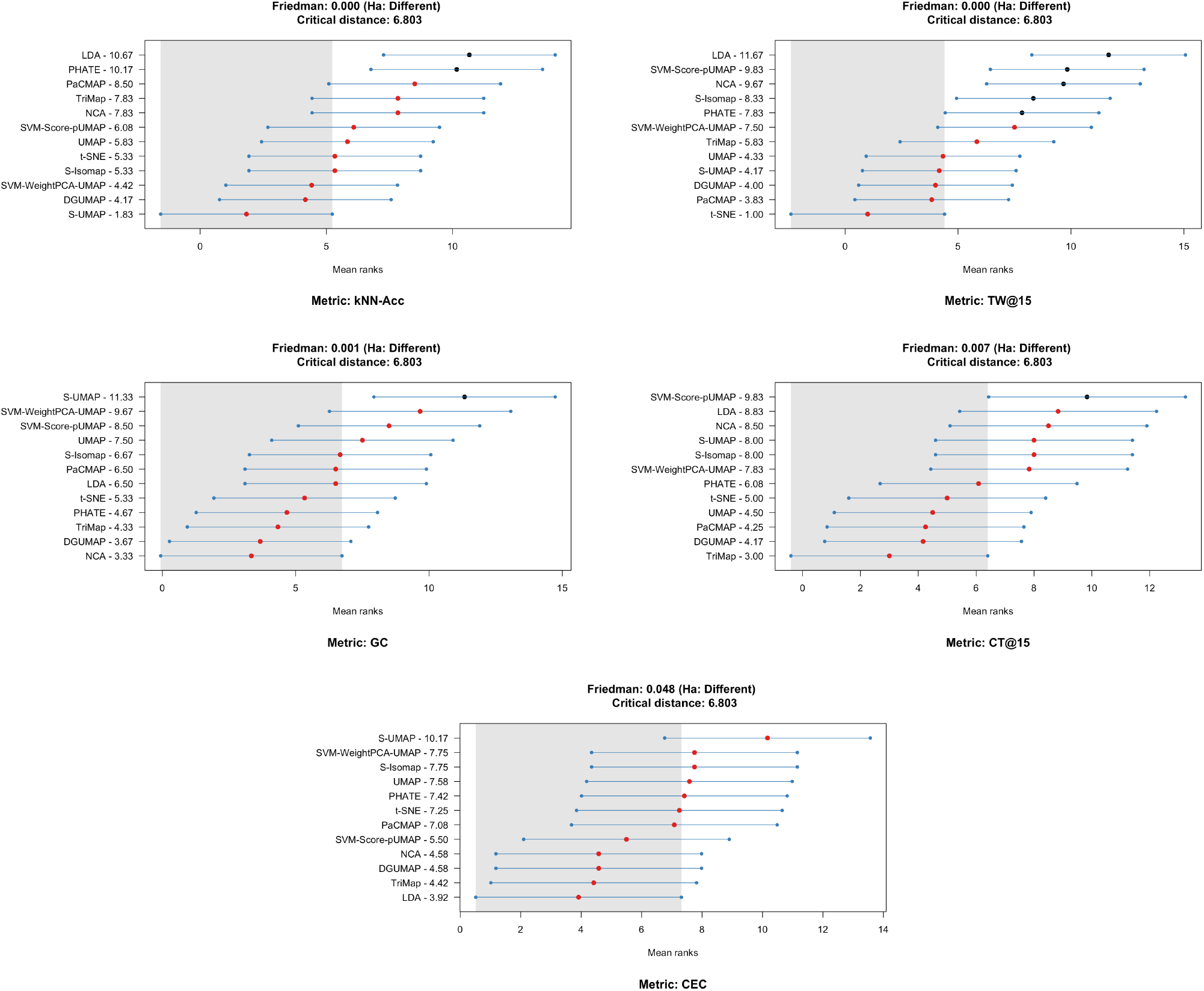
Friedman–Nemenyi rank analysis across datasets for five evaluation metrics. Lower average ranks indicate better performance. Red points show mean ranks, horizontal intervals show Nemenyi critical-distance intervals, and the shaded region denotes the multiple-comparison-with-the-best region at *α* = 0 .05. Methods overlapping the shaded region are not significantly different from the best-ranked method. DG-UMAP consistently falls within this statistically indistinguishable top-performing group, demonstrating robust performance across the evaluated metrics.

## 5 Conclusion

In this work, we introduced DG-UMAP, a supervised dimensionality reduction framework that incorporates label information through decision geometry rather than by directly enforcing class separation in the embedding space. The key idea is to extract boundary-local gradients from a multiclass decision model in the original feature space, aggregate them into a low-rank anisotropic metric, and then apply Parametric UMAP to the resulting transformed representation. In this way, supervision acts by deforming the ambient geometry before embedding, rather than by imposing class-forcing objectives in the low-dimensional space.

Across a diverse set of synthetic and real biomedical datasets, DG-UMAP consistently achieved a strong balance between global geometric faithfulness, local neighborhood preservation, and class-level organization. On structured synthetic datasets such as concentric cylinders and high-dimensional tree manifolds, it more faithfully recovered the expected global arrangement than competing unsupervised and supervised baselines. On real-world datasets including MNIST, single-cell transcriptomic data, Mammoth 3D, and QT-PAD, it yielded embeddings with improved alignment to class structure while avoiding the excessive over-separation often observed in existing supervised methods. These results support the central hypothesis of this work: decision geometry provides a principled and effective mechanism for incorporating supervision without sacrificing manifold faithfulness.

Future work may extend this framework to more general decision models and multimodal biomedical representations. In addition, the polyhedral arrangement induced by multiclass SVM margins may provide a useful prototype global structure for supervised dimensionality reduction, enabling future methods to model class adjacency and higher-order class organization more explicitly than boundary-local geometry alone.

## A A Method Details

### A.1 Random Fourier feature approximation

For the RFF approximation,

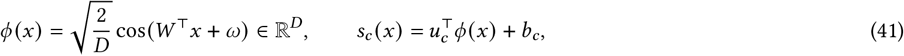

So

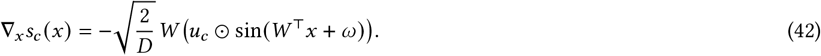

### A.2 Closed form of the target pair distribution

#### Proposition A.1

(Closed form of the target pair distribution). *Let*

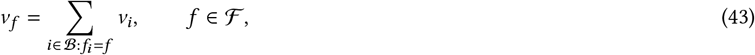

*and let u* ∈ Δ(ℱ) *be any strictly positive reference distribution. Because f* ∈ ℱ *means that there exists i* ∈ B *with f*_*i*_ *i* ∈ ℬ, *we have v*_*f*_ *>* 0 *for every f* ∈ ℱ .

*For β* ∈ [0, 1], *consider*

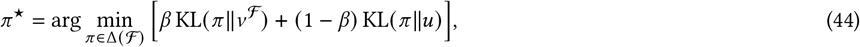

*where v*^ℱ^ = (*v*_*f*_)_*f* ∈_ ℱ. *Then the unique minimizer is*

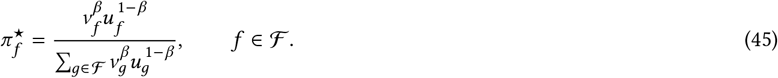

*In particular, if u is uniform on ℱ, i*.*e*.

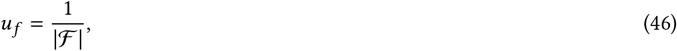

*Then*

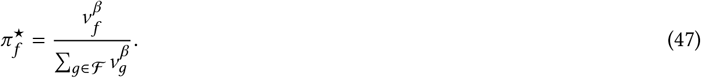

Proof. Define

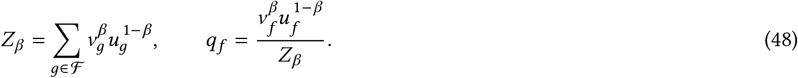

Then *q* ∈ Δ(ℱ). For any *π* ∈ Δ(ℱ),

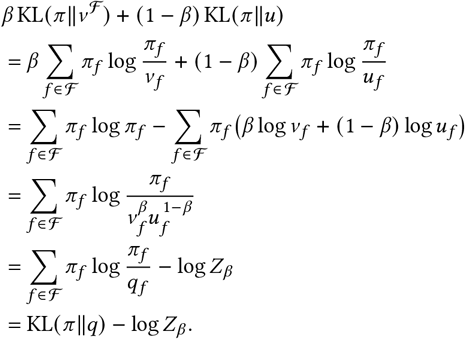

Since KL(*π* ∥*q*) ≥ 0, with equality if and only if *π* = *q*, the unique minimizer is *π*^★^ = *q*. This gives

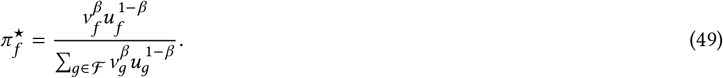

If *u* is uniform, then 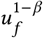 is constant in *f* and is absorbed into the normalization, yielding

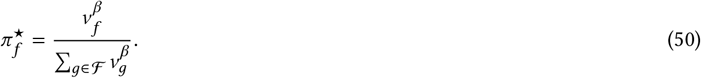

### A.3 Closed form of the sample weights under fixed pair marginals

#### Proposition A.2

(Closed form of the sample weights for fixed pair marginal). *Fix π* ∈ Δ(ℱ) *with π*_*f*_ *>* 0 *for all f* ∈ ℱ . *Consider*

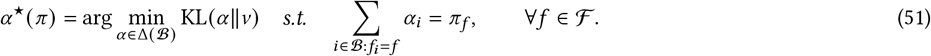

*Then the unique minimizer is*

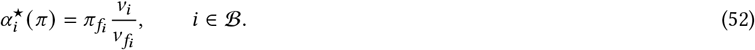

Proof. For each *f* ∈ ℱ, let

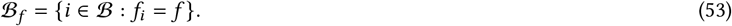

For any feasible *α*, define the conditional distribution within pair *f* by

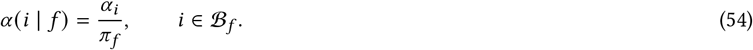

Likewise define the conditional distribution induced by *v*:

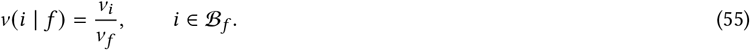

Using *α*_*i*_ = *π*_*f*_ *α* (*i* | *f*) and *v*_*i*_ = *v*_*f*_ *v* (*i* | *f*) for *i* ∈ ℬ_*f*_, we obtain

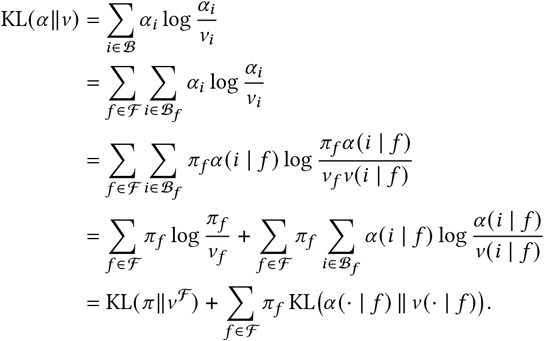

Under the constraint

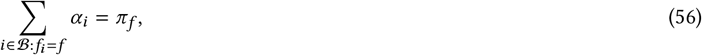

the first term is constant because it depends only on *π* . The second term is nonnegative and equals zero if and only if

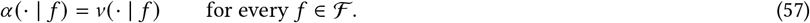

Therefore the unique minimizer is obtained by setting

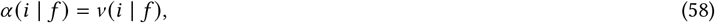

which gives

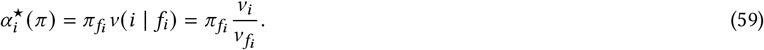

#### Corollary A.3

(Closed form of *α*^★^). *Combining the two propositions yields*

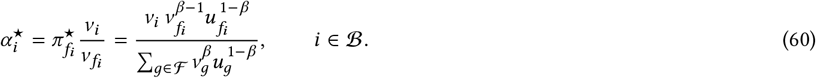

*If u is uniform on* ℱ, *then*

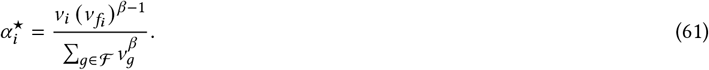

### A.4 Optimal subspace property of *c*

For any rank-*r* orthogonal projector *P*,

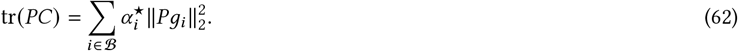

Therefore, by the Ky Fan variational principle, the projector onto the top-*r* eigenspace of *c* maximizes the captured pair-balanced boundary-gradient energy. Equivalently, it minimizes

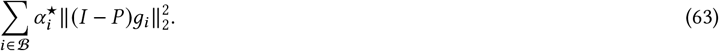

Hence the retained eigenspace is the best rank-*r* subspace for approximating the pair-balanced boundary geometry.

## B Numerical Results

**Table 1:** Performance comparison on the Cylinders_std dataset.

| Method | GC | TW | CT | CEC | kNN-Acc |
| --- | --- | --- | --- | --- | --- |
| DG-UMAP | $0.9348 \pm 0.0032$ | $0.9482 \pm 0.0012$ | $0.9758 \pm 0.0002$ | $0.8478 \pm 0.0050$ | $0.9970 \pm 0.0013$ |
| UMAP | $0.2146 \pm 0.0267$ | $0.9862 \pm 0.0026$ | $0.9903 \pm 0.0033$ | $-0.2873 \pm 0.1606$ | $1.0000 \pm 0.0000$ |
| S-UMAP | $0.2256 \pm 0.0439$ | $0.9922 \pm 0.0024$ | $0.9938 \pm 0.0002$ | $-0.3600 \pm 0.0036$ | $1.0000 \pm 0.0000$ |
| SVM-Score-pUMAP | $0.5464 \pm 0.2050$ | $0.7203 \pm 0.1229$ | $0.9458 \pm 0.0342$ | $0.8381 \pm 0.0093$ | $0.5688 \pm 0.1793$ |
| SVM-WeightPCA-UMAP | $0.2071 \pm 0.0257$ | $0.9877 \pm 0.0034$ | $0.9904 \pm 0.0024$ | $-0.2154 \pm 0.1966$ | $1.0000 \pm 0.0000$ |
| t-SNE | $0.7261 \pm 0.0951$ | $0.9959 \pm 0.0009$ | $0.9709 \pm 0.0012$ | $0.6550 \pm 0.0843$ | $0.9999 \pm 0.0002$ |
| TriMap | $0.6984 \pm 0.1515$ | $0.8731 \pm 0.0480$ | $0.9883 \pm 0.0023$ | $0.8009 \pm 0.0244$ | $0.8795 \pm 0.0555$ |
| PHATE | $0.9275 \pm 0.0081$ | $0.9249 \pm 0.0049$ | $0.9791 \pm 0.0011$ | $0.7895 \pm 0.0328$ | $0.9607 \pm 0.0143$ |
| PaCMAP | $0.4746 \pm 0.0768$ | $0.9339 \pm 0.0236$ | $0.9763 \pm 0.0028$ | $0.2156 \pm 0.4319$ | $0.9047 \pm 0.0489$ |
| NCA | $0.9323 \pm 0.0485$ | $0.9370 \pm 0.0012$ | $0.9817 \pm 0.0023$ | $0.8463 \pm 0.0051$ | $1.0000 \pm 0.0000$ |
| LDA | $0.6815 \pm 0.1538$ | $0.7878 \pm 0.0577$ | $0.9860 \pm 0.0005$ | $0.8455 \pm 0.0052$ | $0.6707 \pm 0.0614$ |
| S-Isomap | $0.2903 \pm 0.0491$ | $0.9560 \pm 0.0075$ | $0.9926 \pm 0.0016$ | $0.3302 \pm 0.0058$ | $1.0000 \pm 0.0000$ |

**Table 2:** Performance comparison on the MNIST_raw784 dataset.

| Method | GC | TW | CT | CEC | kNN-Acc |
| --- | --- | --- | --- | --- | --- |
| DG-UMAP | $0.3213 \pm 0.0320$ | $0.9439 \pm 0.0034$ | $0.9316 \pm 0.0035$ | $0.6662 \pm 0.0492$ | $0.9435 \pm 0.0107$ |
| UMAP | $0.3165 \pm 0.0091$ | $0.9416 \pm 0.0038$ | $0.9344 \pm 0.0025$ | $0.7546 \pm 0.0206$ | $0.9367 \pm 0.0090$ |
| S-UMAP | $0.1862 \pm 0.0543$ | $0.9426 \pm 0.0024$ | $0.8958 \pm 0.0109$ | $0.2564 \pm 0.1907$ | $0.9948 \pm 0.0009$ |
| SVM-Score-pUMAP | $0.2553 \pm 0.0584$ | $0.8438 \pm 0.0066$ | $0.9070 \pm 0.0049$ | $0.7953 \pm 0.0480$ | $0.9345 \pm 0.0045$ |
| SVM-WeightPCA-UMAP | $0.3009 \pm 0.0063$ | $0.8478 \pm 0.0026$ | $0.9116 \pm 0.0031$ | $0.8167 \pm 0.0297$ | $0.9301 \pm 0.0068$ |
| t-SNE | $0.3797 \pm 0.0135$ | $0.9541 \pm 0.0016$ | $0.9370 \pm 0.0029$ | $0.6909 \pm 0.0141$ | $0.9393 \pm 0.0052$ |
| TriMap | $0.1720 \pm 0.0147$ | $0.9298 \pm 0.0036$ | $0.9238 \pm 0.0037$ | $0.8761 \pm 0.0175$ | $0.9312 \pm 0.0105$ |
| PHATE | $0.3815 \pm 0.0173$ | $0.8522 \pm 0.0125$ | $0.9355 \pm 0.0010$ | $0.7609 \pm 0.0152$ | $0.7613 \pm 0.0174$ |
| PaCMAP | $0.2777 \pm 0.0325$ | $0.9397 \pm 0.0030$ | $0.9319 \pm 0.0032$ | $0.6442 \pm 0.0641$ | $0.9026 \pm 0.0159$ |
| NCA | $0.4462 \pm 0.0324$ | $0.7364 \pm 0.0043$ | $0.8765 \pm 0.0024$ | $0.6856 \pm 0.0792$ | $0.7071 \pm 0.0048$ |
| LDA | $0.3229 \pm 0.0250$ | $0.7000 \pm 0.0091$ | $0.8646 \pm 0.0027$ | $0.7764 \pm 0.0200$ | $0.5291 \pm 0.0391$ |
| S-Isomap | $0.5457 \pm 0.0174$ | $0.7631 \pm 0.0076$ | $0.9258 \pm 0.0023$ | $0.5689 \pm 0.0302$ | $0.4902 \pm 0.0204$ |

**Table 3:** Performance comparison on the Mammoth_3D dataset.

| Method | GC | TW | CT | CEC | kNN-Acc |
| --- | --- | --- | --- | --- | --- |
| DG-UMAP | $0.8390 \pm 0.0163$ | $0.9824 \pm 0.0074$ | $0.9957 \pm 0.0005$ | $0.8695 \pm 0.0416$ | $0.9874 \pm 0.0160$ |
| UMAP | $0.7747 \pm 0.0133$ | $0.9798 \pm 0.0048$ | $0.9809 \pm 0.0030$ | $0.8434 \pm 0.0159$ | $0.9870 \pm 0.0022$ |
| S-UMAP | $0.2223 \pm 0.0784$ | $0.9855 \pm 0.0042$ | $0.9516 \pm 0.0083$ | $0.1096 \pm 0.0717$ | $1.0000 \pm 0.0000$ |
| SVM-Score-pUMAP | $0.8402 \pm 0.0346$ | $0.9811 \pm 0.0026$ | $0.9941 \pm 0.0012$ | $0.8904 \pm 0.0332$ | $0.9647 \pm 0.0093$ |
| SVM-WeightPCA-UMAP | $0.7782 \pm 0.0125$ | $0.9788 \pm 0.0040$ | $0.9807 \pm 0.0027$ | $0.8428 \pm 0.0041$ | $0.9888 \pm 0.0023$ |
| t-SNE | $0.5944 \pm 0.0079$ | $0.9940 \pm 0.0005$ | $0.9915 \pm 0.0002$ | $0.5785 \pm 0.0380$ | $0.9857 \pm 0.0009$ |
| TriMap | $0.8669 \pm 0.0029$ | $0.9670 \pm 0.0020$ | $0.9960 \pm 0.0002$ | $0.7484 \pm 0.0239$ | $0.9810 \pm 0.0034$ |
| PHATE | $0.4568 \pm 0.0846$ | $0.9499 \pm 0.0021$ | $0.9913 \pm 0.0004$ | $0.2334 \pm 0.1404$ | $0.9384 \pm 0.0152$ |
| PaCMAP | $0.7943 \pm 0.0101$ | $0.9871 \pm 0.0035$ | $0.9949 \pm 0.0010$ | $0.8254 \pm 0.0313$ | $0.9798 \pm 0.0038$ |
| NCA | $0.8593 \pm 0.0173$ | $0.9484 \pm 0.0076$ | $0.9945 \pm 0.0002$ | $0.8902 \pm 0.0431$ | $0.9181 \pm 0.0086$ |
| LDA | $0.8938 \pm 0.0059$ | $0.9342 \pm 0.0031$ | $0.9946 \pm 0.0001$ | $0.8570 \pm 0.0221$ | $0.8955 \pm 0.0219$ |
| S-Isomap | $0.2866 \pm 0.1532$ | $0.9369 \pm 0.0022$ | $0.9696 \pm 0.0045$ | $0.0218 \pm 0.1219$ | $0.9691 \pm 0.0127$ |

**Table 4:** Performance comparison on the SingleCell_PCA dataset.

| Method | GC | TW | CT | CEC | kNN-Acc |
| --- | --- | --- | --- | --- | --- |
| DG-UMAP | $0.7444 \pm 0.0083$ | $0.9687 \pm 0.0013$ | $0.9741 \pm 0.0011$ | $0.4493 \pm 0.1562$ | $0.9944 \pm 0.0011$ |
| UMAP | $0.5448 \pm 0.0161$ | $0.9673 \pm 0.0011$ | $0.9731 \pm 0.0013$ | $0.0979 \pm 0.0694$ | $0.9815 \pm 0.0006$ |
| S-UMAP | $0.4210 \pm 0.0822$ | $0.9656 \pm 0.0014$ | $0.9668 \pm 0.0029$ | $0.5300 \pm 0.0340$ | $0.9992 \pm 0.0002$ |
| SVM-Score-pUMAP | $0.5313 \pm 0.0349$ | $0.9180 \pm 0.0017$ | $0.9581 \pm 0.0019$ | $0.5921 \pm 0.1399$ | $0.9830 \pm 0.0007$ |
| SVM-WeightPCA-UMAP | $0.5760 \pm 0.0089$ | $0.9355 \pm 0.0036$ | $0.9620 \pm 0.0025$ | $0.1200 \pm 0.0807$ | $0.9811 \pm 0.0008$ |
| t-SNE | $0.6001 \pm 0.0032$ | $0.9724 \pm 0.0009$ | $0.9722 \pm 0.0017$ | $0.4007 \pm 0.0433$ | $0.9839 \pm 0.0011$ |
| TriMap | $0.7718 \pm 0.0100$ | $0.9662 \pm 0.0016$ | $0.9778 \pm 0.0010$ | $0.4786 \pm 0.0662$ | $0.9805 \pm 0.0012$ |
| PHATE | $0.7926 \pm 0.0141$ | $0.9382 \pm 0.0160$ | $0.9713 \pm 0.0047$ | $0.1436 \pm 0.2037$ | $0.9698 \pm 0.0114$ |
| PaCMap | $0.7260 \pm 0.0035$ | $0.9699 \pm 0.0009$ | $0.9770 \pm 0.0012$ | $0.4043 \pm 0.0358$ | $0.9814 \pm 0.0012$ |
| NCA | $0.8469 \pm 0.0033$ | $0.8829 \pm 0.0021$ | $0.9400 \pm 0.0013$ | $0.6314 \pm 0.0257$ | $0.9716 \pm 0.0011$ |
| LDA | $0.4497 \pm 0.0164$ | $0.7629 \pm 0.0020$ | $0.8789 \pm 0.0020$ | $0.5979 \pm 0.0396$ | $0.8883 \pm 0.0019$ |
| S-Isomap | $0.5960 \pm 0.0060$ | $0.9187 \pm 0.0018$ | $0.9366 \pm 0.0011$ | $0.3886 \pm 0.0733$ | $0.9940 \pm 0.0009$ |

**Table 5:** Performance comparison on the PHATE_tree dataset.

| Method | GC | TW | CT | CEC | kNN-Acc |
| --- | --- | --- | --- | --- | --- |
| DG-UMAP | $0.7567 \pm 0.0458$ | $0.9985 \pm 0.0003$ | $0.9985 \pm 0.0002$ | $0.7895 \pm 0.0911$ | $0.9907 \pm 0.0034$ |
| UMAP | $0.7186 \pm 0.0134$ | $0.9982 \pm 0.0007$ | $0.9986 \pm 0.0002$ | $0.7215 \pm 0.0346$ | $0.9821 \pm 0.0071$ |
| S-UMAP | $0.3577 \pm 0.0219$ | $0.9975 \pm 0.0022$ | $0.9887 \pm 0.0013$ | $0.2579 \pm 0.0897$ | $0.9987 \pm 0.0023$ |
| SVM-Score-pUMAP | $0.3814 \pm 0.0588$ | $0.9474 \pm 0.0015$ | $0.9583 \pm 0.0010$ | $0.2229 \pm 0.1454$ | $1.0000 \pm 0.0000$ |
| SVM-WeightPCA-UMAP | $0.3542 \pm 0.0523$ | $0.9718 \pm 0.0009$ | $0.9706 \pm 0.0021$ | $0.1783 \pm 0.0653$ | $1.0000 \pm 0.0000$ |
| t-SNE | $0.8036 \pm 0.0035$ | $0.9993 \pm 0.0000$ | $0.9992 \pm 0.0000$ | $0.6994 \pm 0.0075$ | $0.9843 \pm 0.0049$ |
| TriMap | $0.8194 \pm 0.0063$ | $0.9984 \pm 0.0007$ | $0.9990 \pm 0.0000$ | $0.7972 \pm 0.0152$ | $0.9824 \pm 0.0047$ |
| PHATE | $0.5312 \pm 0.0978$ | $0.9825 \pm 0.0045$ | $0.9979 \pm 0.0002$ | $0.6730 \pm 0.0367$ | $0.9352 \pm 0.0166$ |
| PaCMap | $0.6381 \pm 0.0065$ | $0.9990 \pm 0.0000$ | $0.9979 \pm 0.0001$ | $0.7665 \pm 0.0078$ | $0.9805 \pm 0.0056$ |
| NCA | $0.6428 \pm 0.0017$ | $0.9559 \pm 0.0009$ | $0.9703 \pm 0.0003$ | $0.5894 \pm 0.0057$ | $0.9605 \pm 0.0088$ |
| LDA | $0.6199 \pm 0.0021$ | $0.8784 \pm 0.0026$ | $0.9403 \pm 0.0009$ | $0.7441 \pm 0.0062$ | $0.7747 \pm 0.0091$ |
| S-Isomap | $0.7670 \pm 0.0024$ | $0.9933 \pm 0.0003$ | $0.9930 \pm 0.0001$ | $0.8533 \pm 0.0059$ | $0.9845 \pm 0.0050$ |

**Table 6:** Performance comparison on the ADNI_QT_PAD dataset.

| Method | GC | TW | CT | CEC | kNN-Acc |
| --- | --- | --- | --- | --- | --- |
| DG-UMAP | $0.7363 \pm 0.0094$ | $0.8694 \pm 0.0039$ | $0.9294 \pm 0.0021$ | $1.0000 \pm 0.0000$ | $0.6988 \pm 0.0238$ |
| UMAP | $0.6809 \pm 0.0096$ | $0.9085 \pm 0.0012$ | $0.9267 \pm 0.0004$ | $1.0000 \pm 0.0000$ | $0.6239 \pm 0.0333$ |
| S-UMAP | $0.2896 \pm 0.1090$ | $0.9043 \pm 0.0017$ | $0.7342 \pm 0.0170$ | $-0.2000 \pm 0.6708$ | $0.9960 \pm 0.0040$ |
| SVM-Score-pUMAP | $0.4599 \pm 0.0044$ | $0.6876 \pm 0.0011$ | $0.7153 \pm 0.0003$ | $0.5000 \pm 0.0000$ | $0.8972 \pm 0.0107$ |
| SVM-WeightPCA-UMAP | $0.4418 \pm 0.0218$ | $0.7036 \pm 0.0038$ | $0.7412 \pm 0.0040$ | $1.0000 \pm 0.0000$ | $0.8948 \pm 0.0182$ |
| t-SNE | $0.6481 \pm 0.0100$ | $0.9386 \pm 0.0019$ | $0.9186 \pm 0.0031$ | $1.0000 \pm 0.0000$ | $0.6797 \pm 0.0312$ |
| TriMap | $0.7358 \pm 0.0068$ | $0.9111 \pm 0.0043$ | $0.9357 \pm 0.0008$ | $1.0000 \pm 0.0000$ | $0.6311 \pm 0.0228$ |
| PHATE | $0.7545 \pm 0.0052$ | $0.8416 \pm 0.0116$ | $0.9138 \pm 0.0058$ | $1.0000 \pm 0.0000$ | $0.5865 \pm 0.0248$ |
| PaCMap | $0.7126 \pm 0.0041$ | $0.9064 \pm 0.0047$ | $0.9367 \pm 0.0012$ | $1.0000 \pm 0.0000$ | $0.6120 \pm 0.0265$ |
| NCA | $0.6624 \pm 0.0000$ | $0.7112 \pm 0.0000$ | $0.7755 \pm 0.0000$ | $1.0000 \pm 0.0000$ | $0.9139 \pm 0.0104$ |
| LDA | $0.6448 \pm 0.0000$ | $0.6958 \pm 0.0000$ | $0.7765 \pm 0.0000$ | $1.0000 \pm 0.0000$ | $0.8502 \pm 0.0194$ |
| S-Isomap | $0.5368 \pm 0.0000$ | $0.8127 \pm 0.0000$ | $0.7340 \pm 0.0000$ | $1.0000 \pm 0.0000$ | $1.0000 \pm 0.0000$ |

**Table 7:** SVM settings and corresponding cross-validation accuracy for each dataset.

| Dataset | SVM type | Selected $C$ | CV acc. |
| --- | --- | --- | --- |
| PHATE_tree | kernel | 1.0 | $0.9779 \pm 0.0019$ |
| SingleCell_PCA | kernel | 10.0 | $0.9914 \pm 0.0001$ |
| Cylinders_std | kernel | 1.0 | $1.0000 \pm 0.0000$ |
| ADNI_QT_PAD | kernel | 10.0 | $0.8842 \pm 0.0053$ |
| MNIST_raw784 | linear | 0.1 | $0.9018 \pm 0.0018$ |
| Mammoth_3D | linear | 100.0 | $0.8120 \pm 0.0344$ |

## References

[1] Andrzej Maćkiewicz and Waldemar Ratajczak. Principal components analysis(pca). Computers & Geosciences, 19(3):303–342, 1993.

[2] Laurens Van der Maaten and Geoffrey Hinton. Visualizing data using t-sne. Journal of machine learning research, 9(11), 2008.

[3] Leland McInnes, John Healy, and James Melville. Umap: Uniform manifold approximation and projection for dimension reduction. arXiv preprint 1802.03426, 2018.

[4] Kevin R Moon, David Van Dijk, Zheng Wang, Scott Gigante, Daniel B Burkhardt, William S Chen, Kristina Yim, Antonia van den Elzen, Matthew J Hirn, Ronald R Coifman, et al. Visualizing structure and transitions in high-dimensional biological data. Nature biotechnology, 37(12):1482–1492, 2019.

[5] Suresh Balakrishnama and Aravind Ganapathiraju. Linear discriminant analysis-a brief tutorial. Institute for Signal and information Processing, 18(1998):1–8, 1998.

[6] Xin Geng, De-Chuan Zhan, and Zhi-Hua Zhou. Supervised nonlinear dimension-ality reduction for visualization and classification. IEEE Transactions on Systems, Man, and Cybernetics, Part B (Cybernetics), 35(6):1098–1107, 2005.

[7] Jacob Goldberger, Geoffrey E Hinton, Sam Roweis, and Russ R Salakhutdinov. Neighbourhood components analysis. Advances in neural information processing systems, 17, 2004.

[8] Tim Sainburg, Leland McInnes, and Timothy Q Gentner. Parametric umapembeddings for representation and semisupervised learning. Neural Computation,33(11):2881–2907, 2021.

[9] Ehsan Amid and Manfred K Warmuth. Trimap: Large-scale dimensionality reduction using triplets. arXiv preprint 1910.00204, 2019.

[10] Yingfan Wang, Haiyang Huang, Cynthia Rudin, and Yaron Shaposhnik. Un-derstanding how dimension reduction tools work: an empirical approach to deciphering t-sne, umap, trimap, and pacmap for data visualization. Journal of Machine Learning Research, 22(201):1–73, 2021.

[11] Haiyang Huang, Yingfan Wang, Cynthia Rudin, and Edward P Browne. Towards a comprehensive evaluation of dimension reduction methods for transcriptomic data visualization. Communications biology, 5(1):719, 2022.

[12] Yann LeCun, Corinna Cortes, and CJ Burges. Mnist handwritten digit database. ATT Labs [Online]. Available: http://yann.lecun.com/exdb/mnist, 2, 2010.

[13] Cankun Wang, Diana Acosta, Megan McNutt, Jiang Bian, Anjun Ma, Hongjun Fu, and Qin Ma. A single-cell and spatial rna-seq database for alzheimer’s disease (ssread). Nature Communications, 15(1):4710, 2024.

[14] Janez Demšar. Statistical comparisons of classifiers over multiple data sets. Journal of Machine learning research, 7(Jan):1–30, 2006.

